# LPS core type diversity in the *Escherichia coli* species and associations with phylogeny and virulence gene repertoire

**DOI:** 10.1101/2021.03.22.436425

**Authors:** Sébastien Leclercq, Maxime Branger, David GE Smith, Pierre Germon

## Abstract

*Escherichia coli* is a very versatile species for which diversity has been explored from various perspectives highlighting, for example, phylogenetic groupings, pathovars as well as a wide range of O serotypes. The highly variable O-antigen, the most external part of the lipopolysaccharide component of the outer membrane of *E. coli*, is linked to the innermost lipid A through the core region of LPS of which 5 different structures, denominated K-12, R1, R2, R3 and R4, have been characterized so far. The aim of the present study was to analyze the prevalence of these LPS core types in the *E. coli* species and explore their distribution in the different *E. coli* phylogenetic groups and in relationship with the virulence gene repertoire. Results indicated an uneven distribution of core types between the different phylogroups, with phylogroup A strains being the most diverse in terms of LPS core types while phylogroups B1, D and E strains were dominated by the R3 type and phylogroups B2 and C strains being dominated by the R1 type. Strains carrying the LEE virulence operon were mostly of the R3 type whatever the phylogroup while, within phylogroup B2, strains carrying a K-12 core all belonged to the complex STc131, one of the major clone of extra-intestinal pathogenic *E. coli* (ExPEC) strains. The origin of this uneven distribution is discussed but remains to be explained, as well as the consequences of carrying a specific core type on the physiology of the bacteria.

**IMPACT STATEMENT:** The diversity of the *E. coli* species in terms of type of infections is well known, and its surface O-antigen which shows some correspondence to pathogenicity. This study is the first one to report in detail the diversity and distribution of the LPS core types in the *E. coli* species using whole genome sequences. This distribution of the 5 know core types is analyzed in the different phylogroups along with their association with specific virulence gene repertoire. Results indicate a non-random distribution of LPS core types in the *E. coli s*pecies and, most interestingly, preferential association of certain core types with either some phylogenetic clades or with particular virulence gene repertoires. This work is a first step towards the identification of the origin of this diversity as well as towards the exploration of the physiological properties associated with specific core types.

**DATA SUMMARY:** Complete list of strains with associated data is available in supplementary table S1.

## INTRODUCTION

The *Escherichia coli* species is characterized by the great diversity of the strains it encompasses. This diversity is illustrated, phenotypically, by more than 180 different serogroups of O antigen described as well as by the diversity of commensals and pathovars. Indeed, phenotypic differences translate in different capacities to cause or not infections. Most *E. coli* strains are considered as commensals of warm-blooded animals with an additional reservoir in the environment (1, 2). Still, pathogenic isolates *E. coli* have long been described in different hosts and are responsible for a variety of intestinal and extra-intestinal infections (3–5).

Phylogenetically, *E. coli* strains have been classified in seven major phylogroups termed A, B1, B2, C, D, E and F with an eight recently identified G phylogroup (2, 4, 6). Although not absolute, some preferential associations exist between certain phylogenetic groups and pathovars. As an example, Extraintestinal Pathogenic *E. coli* strains (ExPEC) belong more frequently to the B2 and D phylogroups while entero-hemorrhagic strains (EHEC) most often belong to the E group, but also belong to B1 and A groups (7).

A characteristic that has been so far overlooked in the study of *E. coli* diversity is that of LPS core types. LPS is the major component of the outer leaflet of the outer membrane of *E. coli* (8). The innermost part of LPS is the lipid A, to which is attached the core region of LPS of which 5 different structures have been characterized so far and, at the most external part, the highly variable O-antigen made of repeated polysaccharide subunits.

Reasons to investigate the diversity of LPS core types stems in part from the multiple properties conferred by LPS to the bacteria. The O-antigen of LPS is responsible for the weak permeability of the OM, preventing the entry of a number of toxic compounds such as antibiotics, and thus acts as a protective barrier for bacteria (9). It is also a key determinant in the resistance of bacteria to complement mediated lysis (10, 11).

Being a conserved structure of Gram-negative bacteria, the lipid A part of LPS is one of the major bacterial constituents efficiently recognized by the innate immune system of the host (12). This recognition is mediated by interaction with different host receptors, in particular the TLR4 receptor, and is modulated by the presence of the O-antigen. The core part of LPS has also been shown to regulate the interaction with host cells through the DC-SIGN receptor (13, 14). Nevertheless, this recognition seems to be restricted to the K-12 LPS core type and linked to the presence of specific N-acetyl glucosamine epitopes (13, 14). Although this has not been studied in details, one could expect that other host interaction properties might depend on the type of core LPS.

So far, the distribution of LPS core types has been investigated in a restricted number of reports. The R1 type is the most prevalent of all core types, with R2, R3, R4 and K-12 core types having significantly lower prevalences (15–17).

Using a set of monoclonal antibodies, Gibb and colleagues investigated the distribution of LPS core types in *E. coli* strains from diverse origin. Most strains were R1 type, with a higher percentage of R1 type in isolates from urine, compared to isolates from feces and blood cultures (18). Conversely, Dissanayake *et al.* found no difference in the distribution of LPS core types in commensal and pathogenic strains (16). A potential link between virulence properties and LPS core types is also suggested by the observation that O157:H7 and VTEC strains are in general of the R3 type (19–21). These reports also showed an uneven distribution of LPS core types between phylogroups. While phylogroup B2 and D strains, which encompass a number of ExPEC isolates, were found restricted to R1 types, all four other core types were represented among phylogroup A strains, still with a majority of strains being R1.

Hence, we wished to establish, based on genome sequence analysis, the prevalence of the different LPS core types in the *E. coli* species and, simultaneously, analyze the potential relationships between LPS core types, phylogenetic groups and the virulence genes repertoire.

## MATERIAL AND METHODS

### Choice of strains

A set of 500 genome sequences from 499 *E. coli* strains and 1 *E. albertii* were downloaded from Genbank on 14^th^ October 2018. The *E. coli* genome sequences were randomly chosen among the 13250 sequences available in Genbank for this species when his study was initiated.

### Phylogenetic analysis

A Maximum likelihood phylogenetic tree covering the different strains of our study was obtained by analysis of the pan-genome single-nucleotide polymorphism (SNP) using ksnp3 program (22), itself calling FastTree2 (23). Results from this initial analysis were analyzed to identify groups of clonal strains whose genome sequences were different by less than 10 snps among the snp positions identified in all 499 strains. Only one strain per group were kept which led to a set of 269 *E. coli* strains. A new pan-genome single-nucleotide polymorphism (SNP) analysis was performed using ksnp3 program to produce a Maximum-Likelihood (ML) phylogenetic tree for these non-clonal strains. The strain of *E. albertii* was included as an outgroup. Graphical representation of the ML-tree was performed using the iTOL web server (24).

### In silico LPS core type determination, ECOR and serogroup typing

The type of core LPS was analyzed based on the presence of different alleles of the *waaL* gene as described previously (25). In silico ECOR typing was performed using the ClermonTyping script (26). Serogroup analysis was performed using serotype specific sequences from ECTyper (https://github.com/phac-nml/ecoli_serotyping). The phylogroup determined by the ClermonTyping was, for a few strains, not consistent with the phylogenetic clustering performed by ksnp. In such situations, the ksnp clustering was retained. ST and STc analyses were performed using the MLSTar package (v 0.1.3) using Achtman scheme and STc were allocated based on pubmlst.org database.

### Virulence genes repertoire analysis

*E. coli* virulence proteins were collected from the ecoli_VF_collection (https://github.com/aleimba/ecoli_VF_collection) (27) and a presence/absence matrix was generated as described in Kempf et al. (25). Clustering of strains according to virulence gene repertoire was performed using the R packages pheatmap (1.0.12). Prior to clustering genes based on their presence/absence profiles, we removed genes that were present in more than 90% of strains of each phylogroup. We also excluded from the analysis genes that were both absent in more than 90% of all strains and absent in more than 50% of strains of each phylogroup. From the 1069 genes in the initial panel, only 579 remained after this filtering step.

The optimal number of clusters was determined using the eclust R package. Distance used in clustering analyses was binary (1 for presence, 0 for absence of the corresponding gene) and the method of aggregation was ward.D2. Optimal number of clusters was thus determined to be 43. For each strain, enrichment score of each group of genes was calculated by counting the number of genes from each group present in this strain. Principal component analysis was performed based on the enrichment score of each group of genes using the FactoMiner R package (v2.3).

### Association analyses

For each type of core, a 2×2 contingency table was generated with the number of strains possessing or not the analyzed specific core and the number of strains positive for each virulence gene group. A strain was considered positive for a virulence gene group when its enrichment score was higher than 0.7. The p-value for the probability that each contingency table was a random distribution was calculated by a fisher exact test using stats (v4.0.2) R package. p-values were then adjusted for multiple comparisons by the Bonferroni method. Heatmaps of p-values were generated using the pheatmap package (v1.0.12).

## RESULTS

### Distribution of LPS core types in the different phylogroups

The aim of this study was to investigate the relationship between the different types of LPS core, the phylogenetic group and the carriage of virulence genes in the *E. coli* species.

To this end, we first randomly selected 500 *Escherichia* genomes in Genbank including 499 *E. coli* genomes and 1 *E. albertii* genome. Despite being randomly selected, an initial snp analysis suggested that some strains might be clonal. Because of this potential clonality, before undertaking more in-depth studies, we removed from our dataset strains that could be considered as representative of the same clones and only one strain per such group was kept (see methods). This filtering resulted in a set of 269 strains to which was added the *E. albertii* genome as outgroup.

Distribution of strains among the main *E. coli* phylogroups in this dataset of 269 strains was as follows: 83 phylogroup A strains (30.7 %), 87 B1 strains (32.2 %), 49 B2 strains (18.1%), 10 C strains (3.7 %), 22 D strains (8.1 %), 14 E strains (5.2 %), 4 F strains (1.5 %) (Figure 1). Prevalence of the different LPS core types in this set of *E. coli* strains were 14 (5.2%) for K-12, 105 (38.9%) for R1, 27 (10.0%) for R2, 84 (31.1%) for R3, 35 (13.0%) for R4 (Figure 1). LPS core type could not be attributed for 5 strains (1.9%)

**Figure 1:**
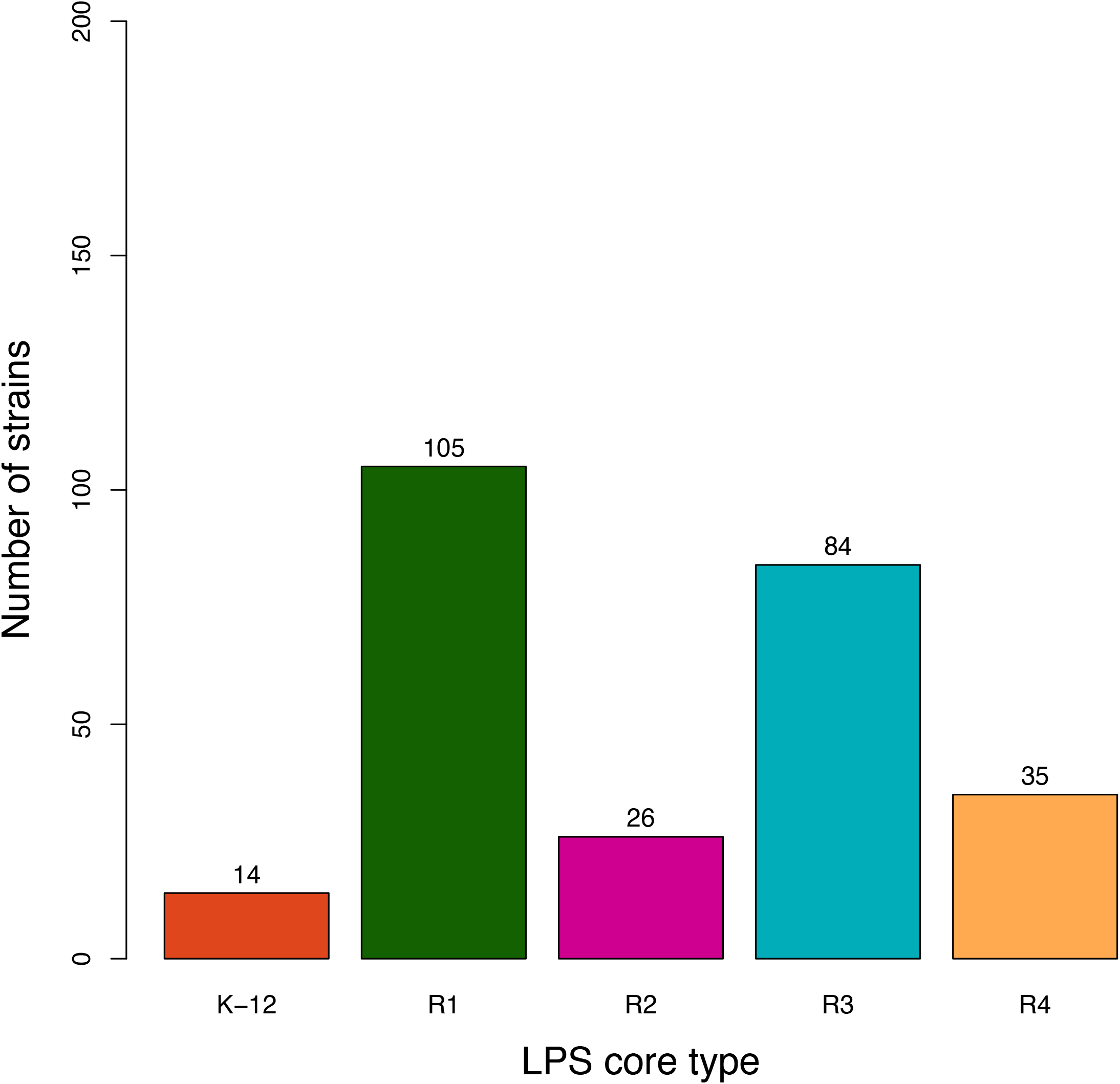
Prevalence of the different LPS core types in *E. coli* strains Core LPS was determined by sequence analysis of core-specific *waaL* alleles. Core type could not be determined for 6 strains.

The prevalence of each core type in each of the different *E. coli* phylogroups was then analyzed which showed that the proportion of the different LPS core types was highly variable depending on the phylogroup of strains (Figure 2). This distribution is non-random as indicated by a Fisher exact test (p<0.001). Phylogoup A strains are very diverse in terms of LPS core types and have somewhat similar proportions of all five LPS core types. R2 strains were mostly found in phylogroup A strains, with two exceptions found in phylogroup E. Strains belonging to the other phylogroups are less diverse with, for example, more strains of the R3 core types in phylogroups B1, D and E. A majority of strains of the B2 phylogroup, as well as all strains of phylogroup C, were of the R1 LPS core type. Finally, the few F strains of our set were either of the R1 or R4 core type.

**Figure 2:**
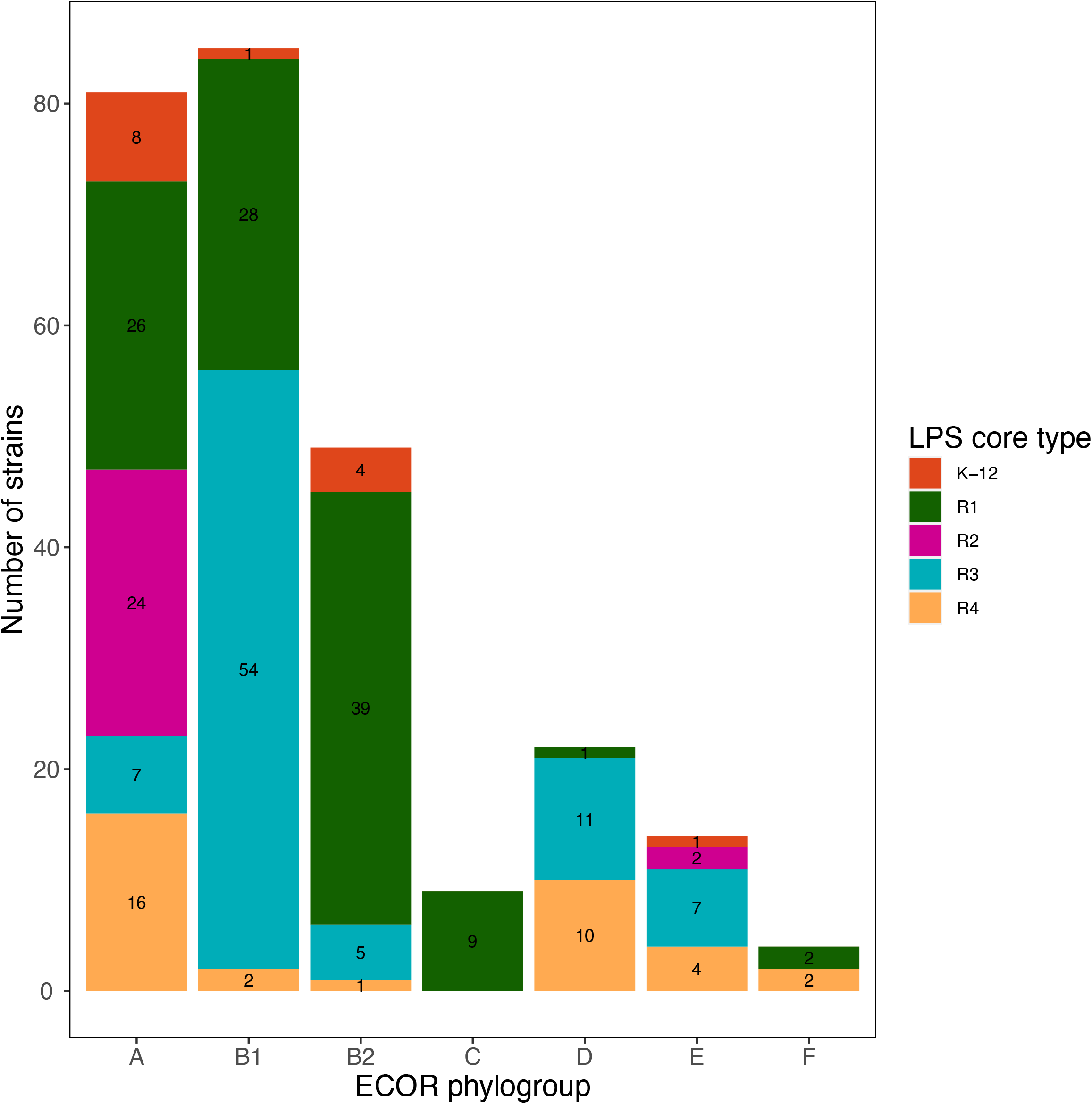
Distribution of LPS core types in major *E. coli* phylogroups. ECOR phylogroup was determined based on ksnp analysis of genome sequences and in silico ECOR typing. Core LPS was determined by sequence analysis of core-specific *waaL* alleles. Only strains belonging to *E. coli* phylogroups A, B1, B2, C, D, E and F are shown.

The serogroup of strains was also determined *in silico*. We observed that some serogroups were preferentially associated with certain LPS core types (Figure 3). For example, strains of serogroup O157 and O73 were preferentially of LPS core type R3 while strains of serogroup O25 were preferentially of LPS core type K-12 or R1. The R1 core type was observed in most O2, O6, O8 and O75 strains. On the contrary, the highly prevalent O1 serogroup showed no preferred association with any LPS core type.

**Figure 3:**
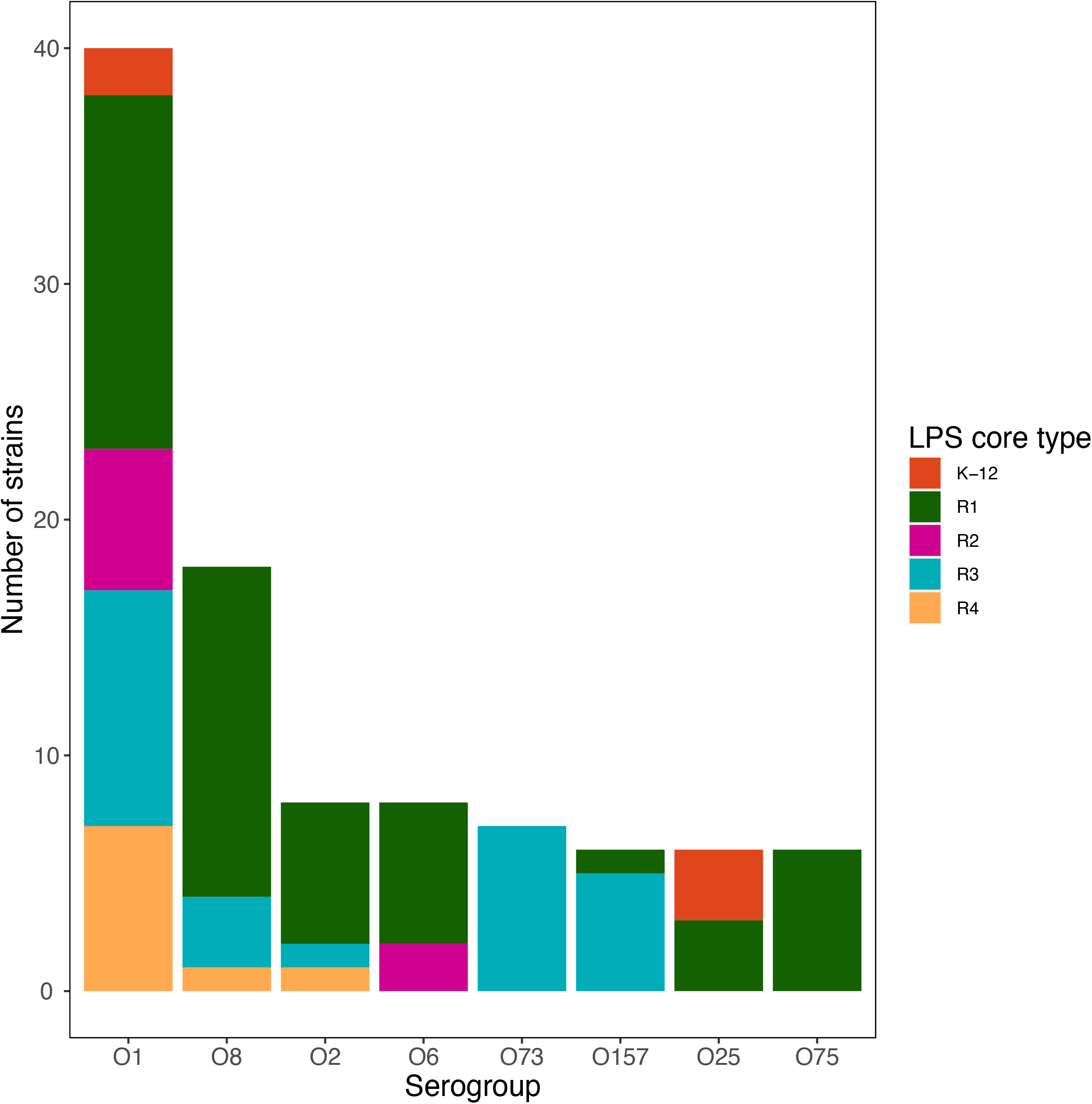
Distribution of LPS core types depending on strains O serogroup. O serogroup was determined based on sequence analyses for O-serogroup specific sequences. Core LPS was determined by sequence analysis of core-specific *waaL* alleles. Only serogroups with more than 10 strains are represented.

### Relationships between phylogeny, LPS core types and virulence gene repertoire

We analyzed the distribution of LPS core types in a genome-wide, SNP-based phylogenetic analysis of the 269 *E. coli* strains. The genome sequence GCA_001286085, annotated as an *E. albertii* sequence in the GenBank database and identified as such by Clermontyper, was used as an outgroup. Again, this analysis clearly differentiated the seven major phylogenetic groups described in the *E. coli* species. The ST complex (STc) to which each strain belongs was determined in silico.

We further investigated the link between phylogeny, specific LPS core types and the carriage of specific virulence genes by defining the repertoire of virulence genes carried by each strain based on the virulence gene dataset published by Leimbach and colleagues [26].

Considering that they would carry less information, we arbitrarily removed genes whose prevalence was either 1) above 90% in strains from each phylogroup or 2) below 30% in all phylogroups, a subset of 596 genes remained.

A hierarchical clustering analysis of virulence genes performed on the 596 remaining genes identified 43 Virulence Gene Groups (VGG) based on their presence/absence patterns (Figure 4). Relevant features of each VGG are presented in Table 1. The complete list of virulence genes in each VGG is presented in supplementary table S2. The hierarchical clustering of VGG was overlayed with the phylogenetic distribution of strains which shows some clearly defined associations of VGG with phylogroups (Figure 4). To better quantify the association between VGG, phylogroups and LPS core types, an enrichment score in genes from each VGG was calculated for each strain and analyzed by principal component analysis (Fig 5A and 5B).

**Figure 4:**
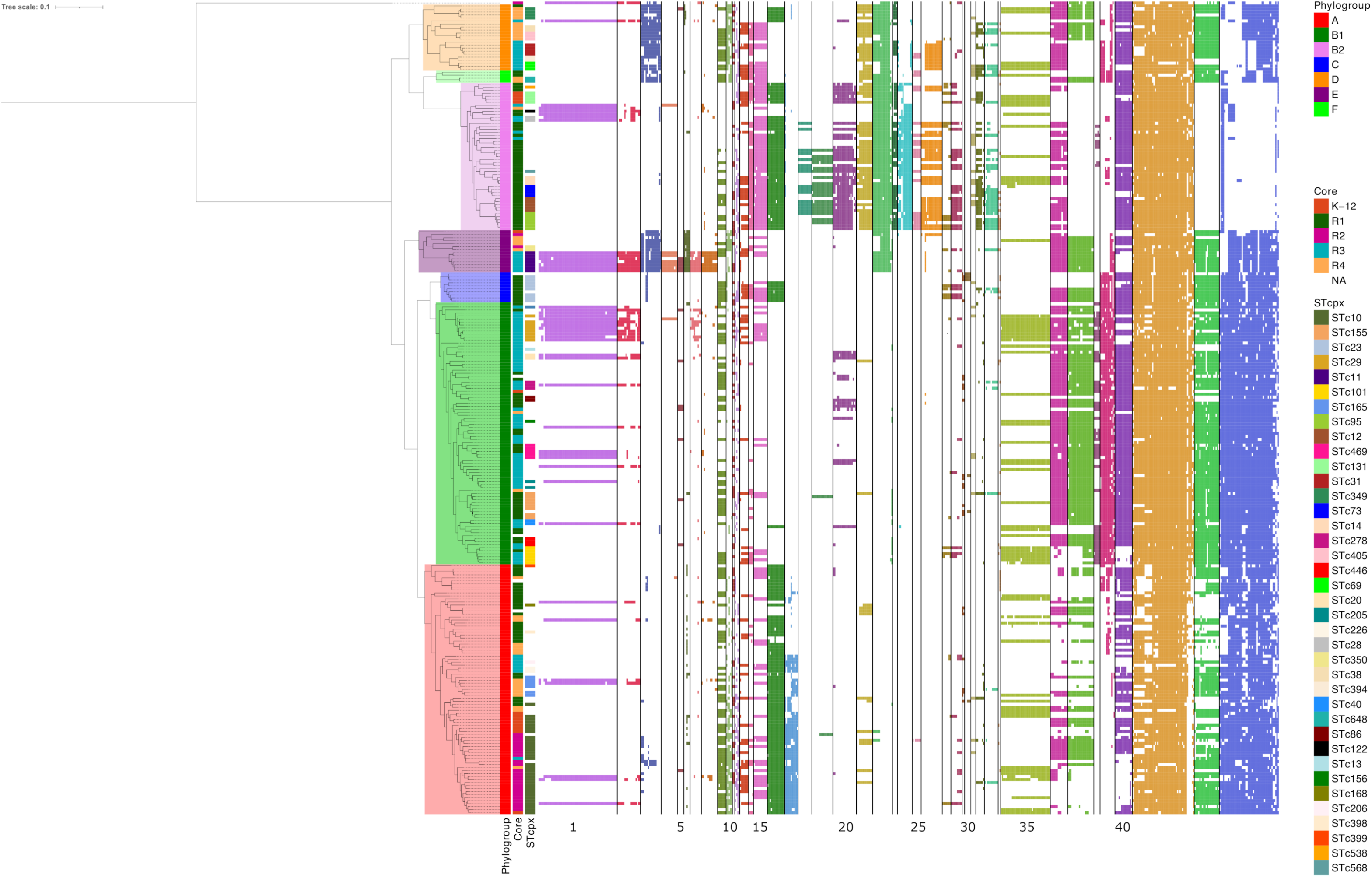
Virulence Gene Repertoire of strains in relation with phylogenetic clustering. Phylogenetic tree of the 269 *E. coli* genome sequences were determined by ksnp analyses. Presence/absence of virulence genes was based on blastp analysis with a threshold of 80% identity and 80% coverage. Presence of a gene is indicated by a colored cell, the color depending on the Virulence Gene groups (VGG) to which the gene belongs. Horizontal lines delineate the different VGG. The strain of *E. albertii* was included as an outgroup. Graphical representation of the ML-tree was performed using the iTOL web server (24).

**Table 1:** Relevant features of significant Virulence Gene Groups

| Group ID | Relevant features |
| --- | --- |
| Group 1 | LEE genes and T3SS effectors |
| Group 2 | LEE genes and T3SS effectors |
| Group 4 | T2SS from EDL933 (*etp* cluster) |
| Group 5 | Shiga-toxin genes |
| Group 6 | *fbp* iron acquisition system |
| Group 7 | T3SS effectors |
| Group 8 | LEE genes and T3SS effectors |
| Group 9 | *fec* iron uptake system |
| Group 13 | Aerobactin iron uptake system |
| Group 14 | *sitABCD* iron uptake system |
| Group 15 | HPI |
| Group 18 | *sfa* and *foc* fimbrial clusters |
| Group 19 | Colibactin operon |
| Group 20 | T6SS type i2 from *E. coli* 536 |
| Group 21 | *kps* gene cluster |
| Group 22 | *chu* and *fit* iron uptake systems |
| Group 24 | *auf* fimbrial cluster |
| Group 25 | K1 capsule cluster |
| Group 26 | T6SS type i1 from *E. coli* APEC O1 |
| Group 28 | *iro* iron uptake system |
| Group 33 | P fimbriae gene cluster |
| Group 35 | Lateral flagella |
| Group 37 | T6SS type i2 from *E. coli* ECC-1470 and EDL933 |
| Group 38 | *lpf* cluster #1 from *E. coli* ATCC9637 |
| Group 39 | *lpf* cluster #2 from *E. coli* ATCC9637 |
| Group 40 | *gsp* T2SS from *E. coli* ATCC9637 |
| Group 41 | Type 1 fimbriae, flagella, *csgAD* curli, *fhuA* and *cirA* iron uptake systems |
| Group 43 | *sfm* fimbriae, *gfc* serum resistance cluster |

**Figure 5:**
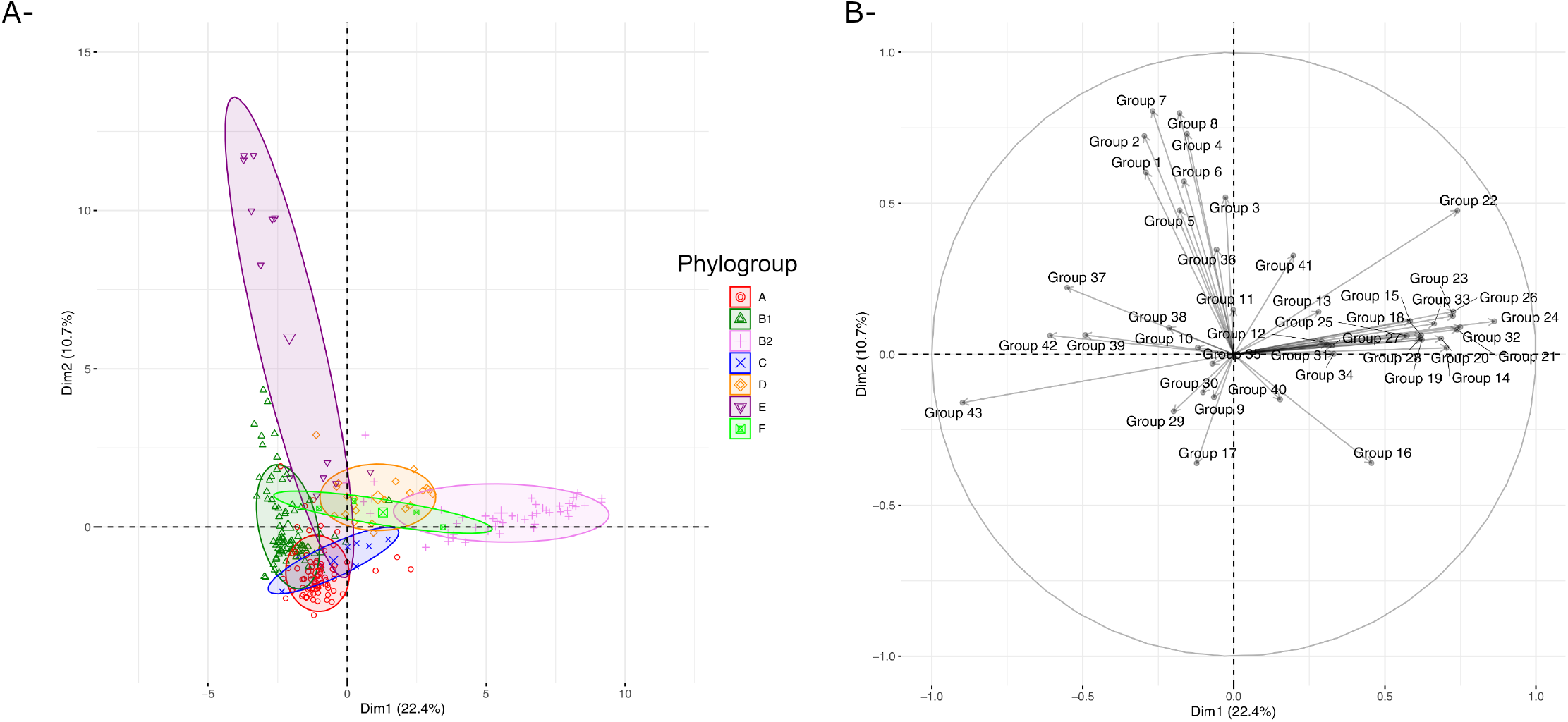
Principal component analysis to analyze the relationships between phylogroups and enrichment in virulence genes. Enrichment of each strain in the different gene groups was calculated by counting the number of virulence genes from a specific VGG in each strain. PCA was performed on the Enrichment scores values using the FactoMineR package. Strains are colored by phylogroup (A) and contribution of the different VGG is indicated (B).

A clear clustering of strains based on phylogroup was observed, in particular for phylogroup B2 strains which were located on the positive side of the Dim1 axis. Some overlap was observed for A, B1, and C strains, based mostly on VGG #9, 17, 29, 30 while B1 strains being mostly positive for VGG #37-39 (T6SS and *lpf* gene clusters). Phylogroup E strains were enriched in VGG #1-8 which include most Type 3 Secretion System (T3SS) structural and effector genes. B2 strains were enriched in VGG #14-33 (iron acquisition genes, HPI, colibactin cluster, *kps* genes, K1 capsule cluster, P fimbriae genes) and were mostly devoid of VGG #1-8, VGG #37-39 and VGG#42-43 genes.

We then overlaid the type of core LPS onto the PCA plot to detect potential associations between VGG and LPS core types. As shown in Figure 6, no well-defined clusters were distinguished when considering all strains together (Figure 6A). When strains for specific phylogroups were analyzed separately some clustering was observed. For example, phylogroup E shows a clear difference in virulence gene content between strains of LPS core type R3 and the others, driven by the presence of VGG #1-6 in R3 types only (Figure 6F). More generally, this finding can be extended to all strains carrying the LEE (VGG #1): LEE positive strains are preferentially of the R3 core type compared to LEE negative strains and this difference is significant (Fisher exact test; p<0.001): out of the 42 strains carrying the LEE and distributed in the different phylogroups, 30 have an R3 core type.

**Figure 6:**
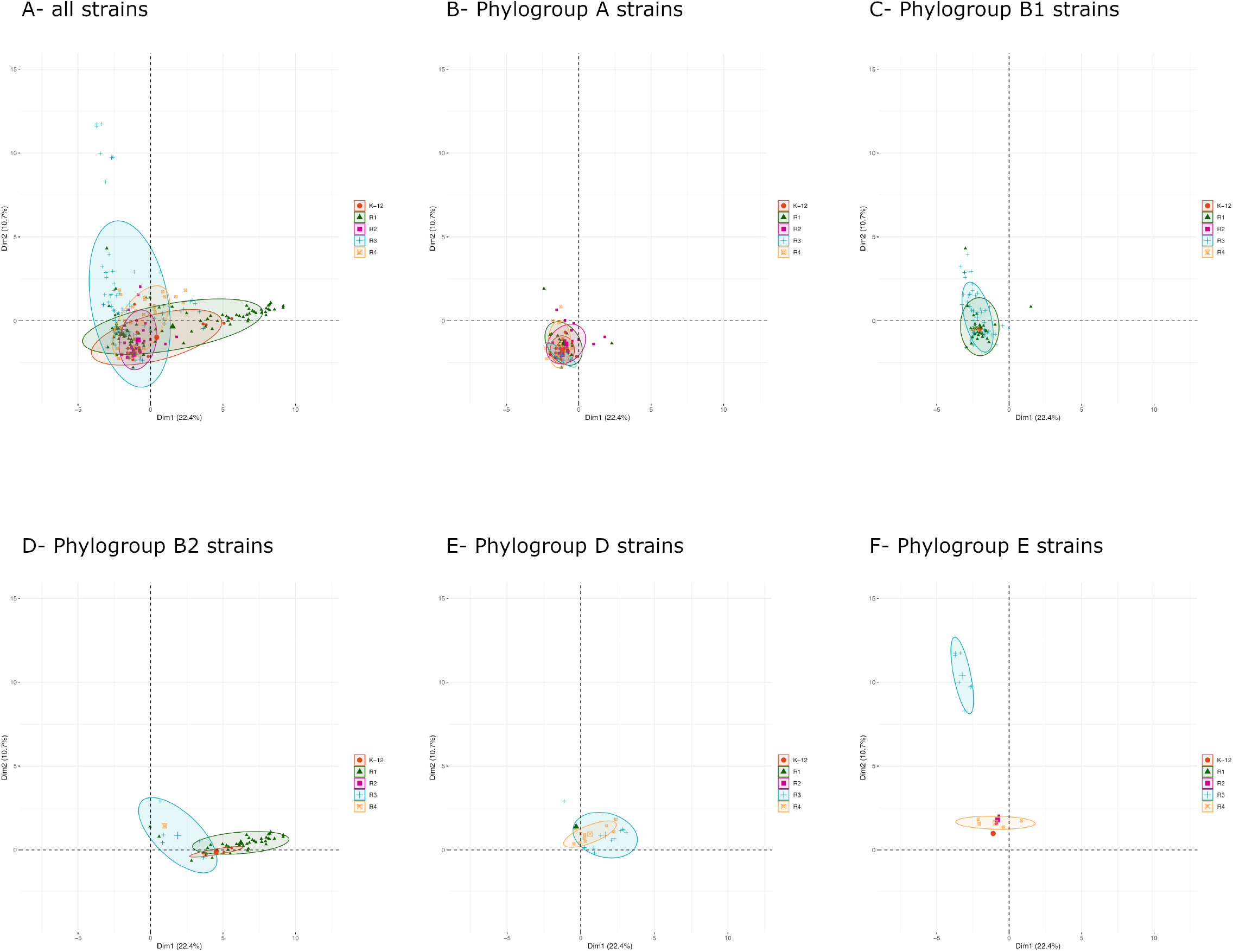
Principal component analysis to analyze the relationships between phylogroups and LPS core types. Enrichment of each strain in the different gene groups was calculated by counting the number of virulence genes from a specific VGG in each strain. PCA was performed on the Enrichment scores values using the FactoMineR package. (A) Strains belonging to all phylogroups are represented. (B-F) Strains belonging to phylogroups A, B1, B2, D and E are represented.

Yet, the most prominent clustering was detected with phylogroup B2 strains. Indeed, the strain distribution allowed the distinction of three clear clusters, one for each LPS core types K-12, R1 and R3 (Figure 6D). Interestingly, B2 strains with a K-12 LPS core all belonged to the STc131 complex. When the 500 strains from our initial dataset were considered, this finding was confirmed (Figure 7). This cluster of B2 strains with a K-12 LPS core types, all belonging to STc131, are enriched in VGG #9, 13 and 35 containing the *fec* operon, the lateral flagellar genes and the aerobactin operon respectively (Fig 7). A group of B2 strains related to the STc28 with a R3 LPS core type are the only ones carrying the VGG#1-2 genes corresponding to the T3SS and some of its effectors. These strains also possess an O-antigen capsule cluster (*etk*, *etp* and *gfcBCDE* genes – found in VGG #43) and are devoid of any of the five T6SS analyzed.

**Figure 7:**
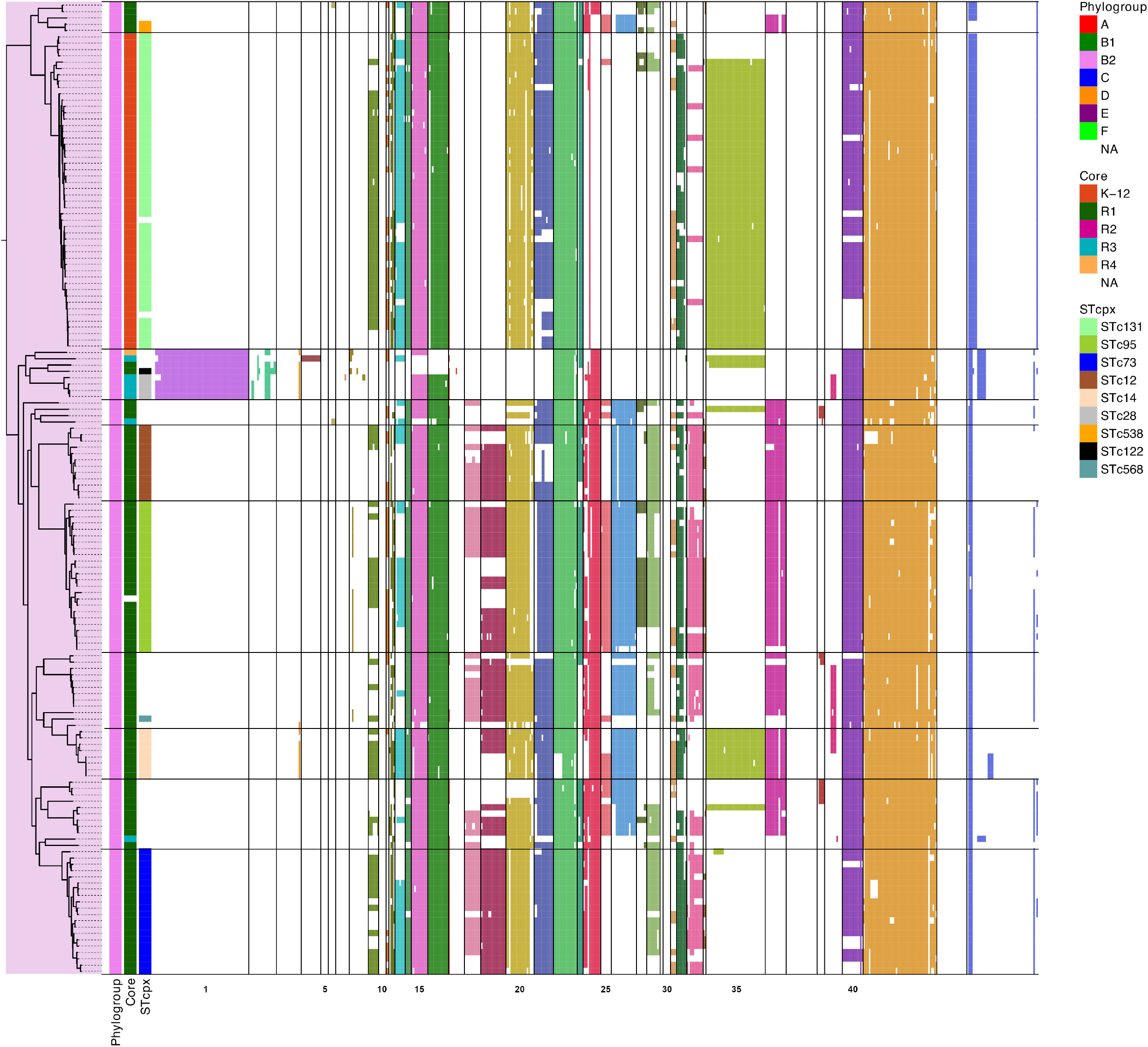
Virulence Gene Repertoire of strains in relation with phylogenetic clustering of phylogroup B2 strains. All B2 strains from the initial 500 genome dataset are represented. Horizontal lines delineate the different B2 STc identified.

Other B2 strains, belonging to the STc73, STc95, STc12 and STc14, almost all carried an R1 core type and were enriched in VGG #18-19, 24-26, 28, 33, and 36 corresponding, among others, to the colibacin operon and several frimbriae gene clusters (Figure 7).

### Distribution of T6SS and iron acquisition genes

Among the virulence genes analyzed, the distribution of Type 6 secretion systems and iron acquisition systems highlighted features of particular interest. First, our virulence dataset covered five different T6SS operons which were not distributed evenly between strains of the different phylogroups (Figure 8). The T6SS from strain 536, belonging to VGG #20, was mostly present in B2 strains, while the one from strain Ecc-1470 (VGG #37) was mostly present in E and B1 strains and absent from B2 strains. The T6SS from strain APEC-O1 (VGG #26) was largely present in B2 strains, with the notable exception of strains from STc131 and STc73 complexes. Interestingly, B2 strains from STc28 did not possess any of the five T6SS. Among D strains, strains possessed either T6SS from APEC-O1 or from ECC-1470 but never both. The two T6SS from strain 042 were the least frequent (and thus are not depicted in Figure 4 nor listed in supplementary Table 2).

**Figure 8:**
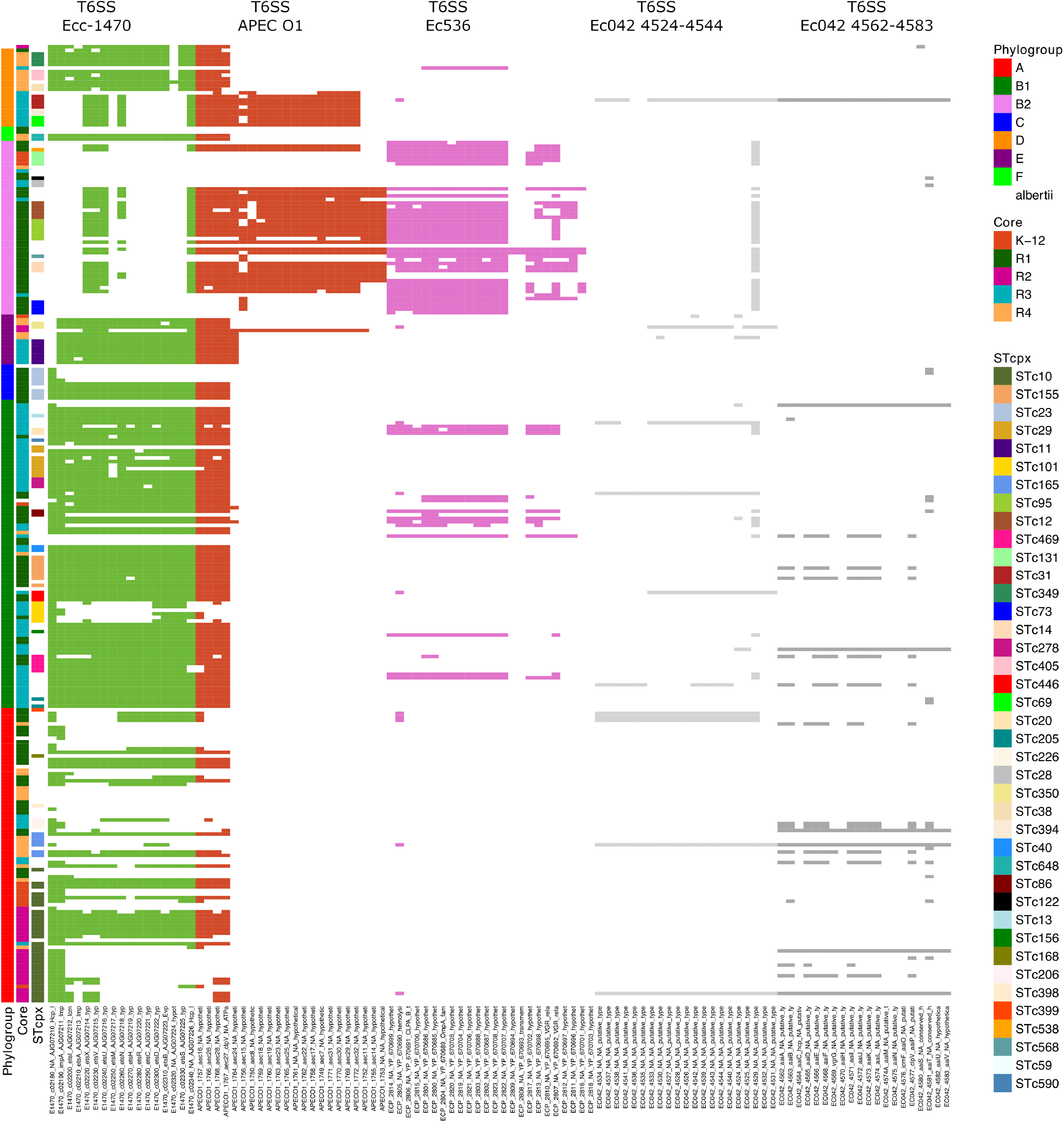
Distribution of Type 6 Secretion Systems in the *E. coli* species. Strains are ordered as in Figure 4. Presence/absence of virulence genes was based on blastp analysis with a threshold of 80% identity and 80% coverage. Presence of genes of the five different T6SS is indicated by a colored cell, the color depending on the T6SS to which the gene belongs.

Concerning iron acquisition systems, our analysis confirmed that some are ubiquitous, such as *fep*, *feo* and *ent* operons, while others are restricted to certain phylogroups (Figure 9). In particular, phylogroup E strains all possessed the *fbpABC* locus while they lacked the yersiniabactin, aerobactin, ferric citrate and *sitABCD* operons. Consistent with the Clermont typing scheme, strains from phylogroup A and B1 always lacked the *chu* operon.

**Figure 9:**
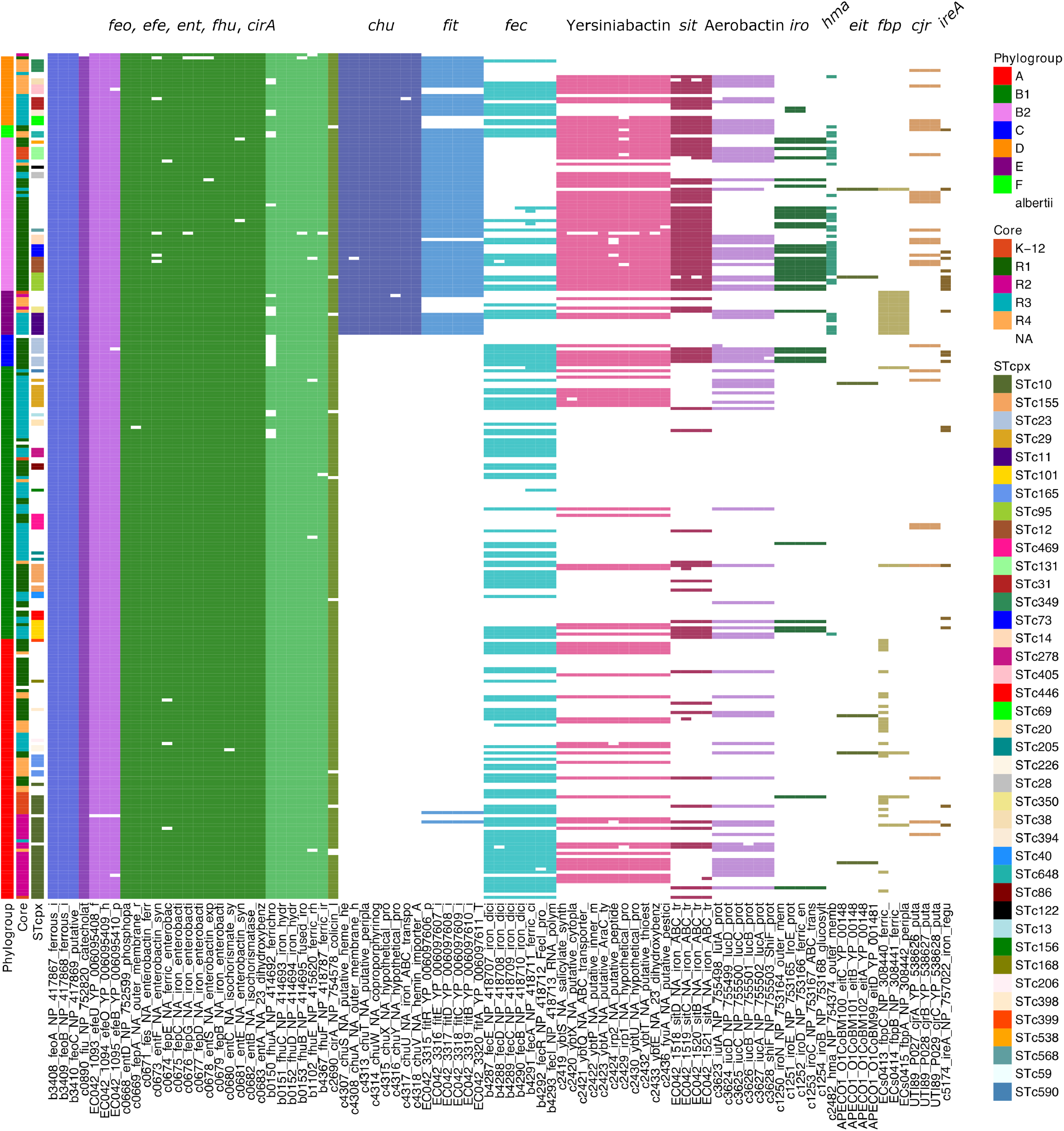
Distribution of iron acquisition systems in the *E. coli* species. Strains are ordered as in Figure 4. Presence/absence of all the iron uptake genes studied in this report was based on blastp analysis with a threshold of 80% identity and 80% coverage.

## DISCUSSION

The initial purpose of this study was to investigate the distribution of the different LPS core types in the *E. coli* species. LPS core types are encoded by the *waa* locus genes located approximately 100kb from the origin of replication in the *E. coli* species. The five distinct regions share highly similar 5’ and 3’ regions, encompassing *waaDFC* and *waaA-waaQGP* genes, while showing variable regions in between with lower GC% values which may correlate to requirement for higher level of transcription for this essential biosynthetic locus (Supplementary figure 1) (28).

After selecting a random set of 500 *E. coli* genomes from Genbank and restricting our dataset to 269 *E. coli* genomes with more than 10 snp differences, we thus set out to examine the prevalence of the difference core types, each core type being characterized by the presence of the O-antigen *waaL* specific ligase as previously described (25).

The high prevalence of the R1 core type we observed had already been described in previous studies analyzing avian pathogenic *E. coli* strains or clinical and commensal isolates (15–17). Another finding consistent with previous reports is the low prevalence of R2 and R4 core types. The high prevalence of R3 core types we observed might be due to an overrepresentation of EHEC/STEC epidemic strains belonging to phylogroup E, as has been observed previously (see below).

The initial phase analyzing the distribution of LPS core types among *E. coli* phylogroups clearly showed a non-random partitioning. As expected, we were able to distinguish the 7 major phylogenetic groups of the *E. coli* species, A, B1, B2, C, D, E and F. This clustering was confirmed by in silico phylogroup determination based on the scheme by Clermont et al. (26). An initial observation was an almost even representation of all five core types among phylogroup A strains while other phylogroups were enriched in some specific core types, in particular E and B1 strains in R3 core types and B2 strains in R1 types. This high diversity observed in phylogroup A strains is reminiscent to the high diversity of gene families and large pan-genome of phylogroup A strains recently described by Touchon *et al.* (29).

This uneven distribution of LPS core types among phylogroups was also observed with serogroups. While only the most prevalent serogroups were analyzed, we found that only the O1 O-antigen could be associated with any of the 5 LPS core types. Conversely, the O157, O73 or O103 O-antigens were preferentially attached to the R3 core type.

A more detailed analysis showed that, within each phylogroup, phylogenetic clusters of strains were observed with specific LPS core types. This is notably true for specific clonal complexes of phylogroup B2 strains where the STc131 complex is associated with K-12 LPS core types and STc28 with R3 core types while most other B2 strains are of the R1 type. Phylogroup D strains also show cluster correlation with specific LPS core types, with R4 and R3 strains belonging to distinct clades in the D phylogroup. Clustering was also obvious in phylogroup E strains with two phylogenetic clades encompassing either strains with an R3 core type and or with R4 types.

One question that justified the present study was whether we could identify a link between specific LPS core types and virulence properties or adaptation to specific environmental niches.

When virulence gene repertoires were analyzed, we could identify co-presence patterns by distinguishing different virulence gene groups. Based on virulence gene repertoire data, clear clustering based on phylogroup was observed confirming that strains belonging to the same phylogenetic group are enriched in some virulence gene sets, consistent with previous studies (29, 30).

Concerning the link between the virulence gene repertoires and the LPS core type, a significant association of the R3 core types with LEE (VGG #1) positive strains was observed, although LEE positive strains with other LPS core types were identified. One hypothesis that could explain such a bias would be that acquisition of the LEE locus preferentially required an R3 core. Indeed, acquisition of the LEE locus through phage transfer has been suggested and one could suggest that such transducing phages could require an R3 core as a receptor (31, 32). Curiously, none of LEE-positive phylogroup A strains possessed an R3 core but rather possessed R1, R2 or R4 core types in roughly equal proportions. Whether this represents distinct acquisition events is an open issue.

Despite this lack of recognizable association between virulence gene repertoire and LPS core types, we further deciphered the diversity of B2 strains and possible relationships between their virulence gene repertoire, LPS core types and serogroup. Another finding of the present work is the specific association of the core LPS K12 with STc131 within phylogroup B2, whatever the serogroup (O1, O25, or O16). STc131 clonal complex is one of the four major ExPEC clones that have been identified in various continents and show frequent resistance to multiple drugs (4). The reason for this strong association between STc131 strains and K-12 core types would deserve further investigation to understand if the K-12 core type of STc131 strains could contribute to the specific properties of these strains, in particular fitness or antibiotic resistance through altered outer membrane permeability for example, or to the emergence of these strains.

More generally, a question that remains unanswered is whether specific LPS core types could be associated with certain ecological niches. In addition to the association of STc131 with K-12 core type, our results show unequal distribution of all five core types observed in the different phylogroups. Above all, the reasons and consequences for some of the above-mentioned associations remain to be decrypted. Different core LPS may confer different properties to the outer membrane of bacteria. It may for instance provide a better ability to interact with different structures such as phage tails or immune cell receptors. Indeed, part of the LPS molecules have been identified as receptors for P1 phage (33). Different LPS core types could thus allow bacteria to escape the binding of these bacteriophages or, conversely, permit phage binding. When one considers the strong selective pressure imposed by bacteriophages on bacterial evolution, possessing a specific LPS core type could be a potential selective advantage in environment loaded with bacteriophages not binding this core type. The core type could also influence how bacteria are recognized by host immune cells, as was shown for K-12 core types interacting with the DC-SIGN receptor at the surface of dendritic cells (13, 14).

A bystander observation of our study was that T6SS distribution varied between strains of different phylogroups. T6SS have recently been shown to contribute to the adaptation of strains to their niche such as the intestinal microbiota. T6SS could thus be a major player for adaptation to different niches, each being colonized by different microbiota (34). To that end, it is interesting to note that, based on strains isolated from different sources in Australia, Touchon *et al.* showed strong associations between the phylogenetic structure of populations and the natural habitats of strains (29). Hence, the interplay between T6SS, phylogenetic group and adaptation to a specific niche will be worth investigating in more details.

Likewise, we observed a clear pattern of iron acquisition genes within several phylogenetic groups. Such observations were previously made for EHEC and UPEC strains in which the *fec* iron transport system was absent or for B2 strains in which the *chuA* gene is absent and serves as a base for the Clermont typing scheme (35).

Altogether, this report highlighted the diversity of LPS core types in the *E. coli* species with specific associations between LPS core types and strains of certain phylogroups. Based on the clustering of some LPS types with specific groupings, such as K-12 core types and STc131 strains, our results support the relevance of analyses of the biological properties of LPS molecules belonging to different LPS core types. This reveals the need for further genotype-phenotype investigations targeting surface structures such as LPS core and ability to occupy distinct niches.

## AUTHORS AND CONTRIBUTORS

PG contributed to conceptualization, data curation, formal analysis, funding acquisition, methodology, project administration, supervision, validation, visualization, writing of the original draft, review and editing.

SL contributed to conceptualization, investigation, methodology, validation, review and editing of the manuscript.

DGES contributed to conceptualization, investigation, methodology, validation, review and editing of the manuscript.

MB contributed to data curation.

