## Supplementary figures and images for "LPS core type diversity in the *Escherichia coli* species and associations with phylogeny and virulence gene repertoire"

### Supplemental Figure 1

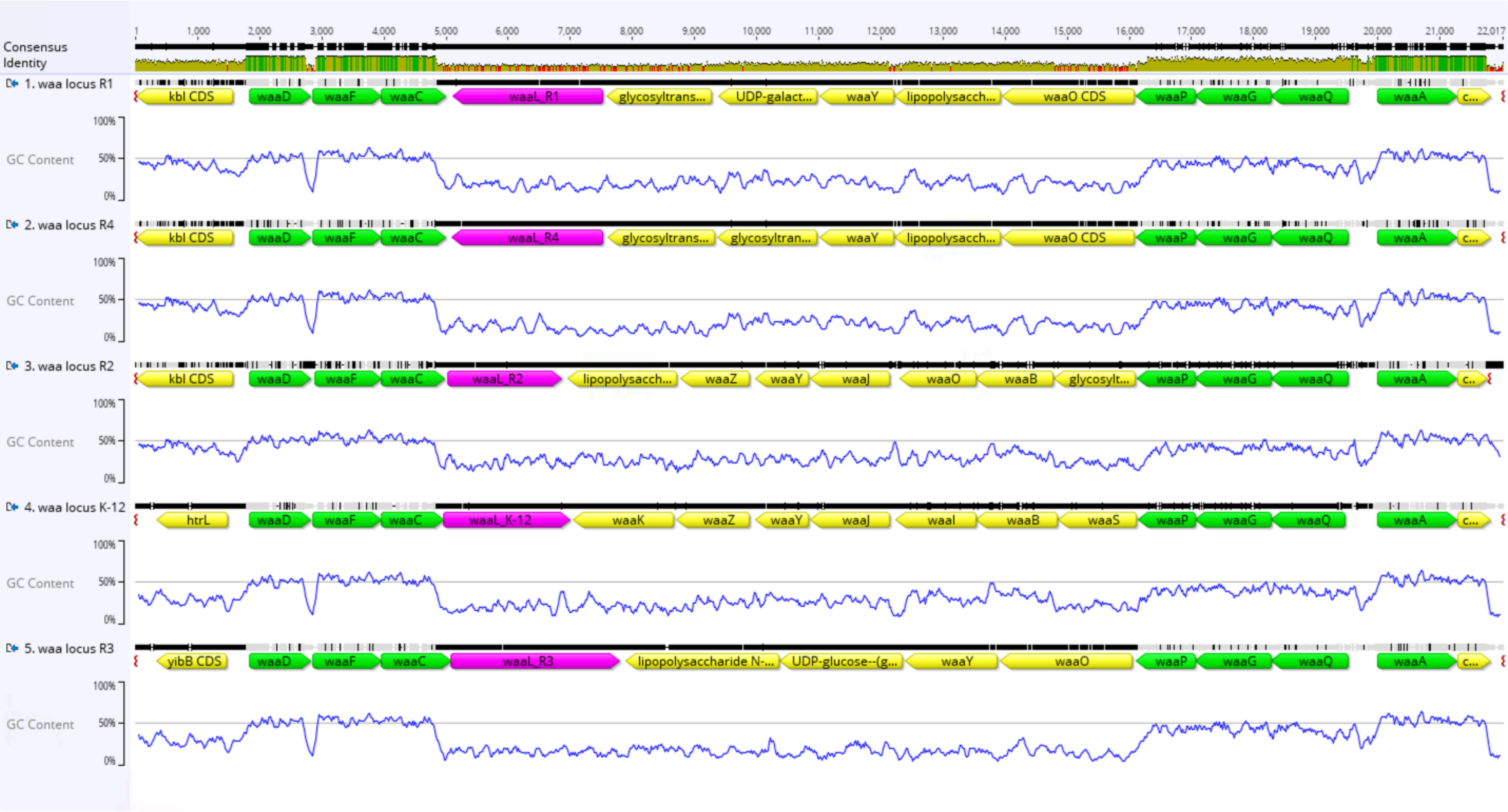
